# No more than three PlpE non-overlapping epitopes trigger significant antibody production in individuals vaccinated with the *Pasteurella multocida* epitope-chimeric proteins

**DOI:** 10.1101/2025.09.02.673733

**Authors:** Binbin Geng, Banghui Zhou, Guojun Jiang, Weifeng Zhu

**Affiliations:** College of Veterinary Medicine, Hebei Agricultural University, Baoding Hebei, 071000, China; Veterinary Biological Technology Innovation Center of Hebei Province, Baoding Hebei 071000, China

## Abstract

Current vaccine research still confronts multiple challenges, and epitope-focused vaccine design serves as an effective technical approach to address these issues. Antigens harbor multiple epitopes; however, the number of epitopes capable of triggering significant antibody responses in vaccinated individuals remains undefined. This study aimed to determine the number of non-overlapping epitopes—designated “effective epitopes” hereafter—that trigger significant antibody production in vaccinated individuals when presented in distinct PlpE epitope-chimeric proteins. Herein, chimeric proteins incorporating varying numbers of PlpE non-overlapping B-cell epitopes were generated. By analyzing serum antibody responses in vaccinated individuals, the number characteristic of PlpE effective epitopes was elucidated. The total number of PlpE effective epitopes in vaccinated individuals was 2.6 (95%CI: 1.909-3.201) for full-length PlpE, and 1.2 (95%CI: 0.5426-1.857), 0.9 (95%CI: 0-1.820), and 0.2 (95%CI: 0-0.5016) for the PlpE chimeric proteins (PlpE-VP60P, PlpE-BcfA, PlpE-PtfA), respectively. With an increase in the number of PlpE non-overlapping epitopes incorporated into the chimeric proteins, the number of PlpE effective epitopes exhibited a trend of initial increase followed by a decrease. Ultimately, the average total number of effective epitopes across all PlpE chimeric proteins did not exceed 3, with the highest number 2.1 (95%CI: 1.572-2.628). In conclusion, the number of PlpE non-overlapping epitopes on an antigen, that trigger significant antibody responses in each vaccinated individual, is very limited.

**IMPORTANCE:** Epitopes underpin the antigenicity of protein antigens. Although the concept of antigenic epitopes has been proposed over 50 years, our understanding on epitopes remains incomplete. Multiple antigenic epitopes can be identified on a single antigen, while the number of these epitopes that function in vaccinated individuals remains unclear—a gap hindering the rational design of vaccines. In previous studies, we identified 6 non-overlapping epitopes of *Pasteurella multocida* PlpE. Herein, we found that the total number of non-overlapping epitopes—capable of significantly triggering antibody production—that are present in PlpE chimeric proteins does not exceed 3 per vaccinated individual. This finding offers important insights for rational vaccine design: given the highly limited number of non-overlapping epitopes that function in vaccinated individuals, only a limited number of epitopes can be grafted onto scaffold proteins. Epitope-focused vaccine design must therefore account for competitive interactions between epitopes on the new antigen.

## OBSERVATION

In both human and veterinary medicine, vaccines play an indispensable role in controlling infectious diseases, effectively reducing the morbidity and mortality associated with these illnesses. Nevertheless, current vaccine research continues to confront challenges, such as the scarcity of effective protective antigens (1), high pathogen variability (2), antibody-dependent enhancement of infection (3), and co-infection pressure (4). Genetic engineering vaccines enable rational vaccine design, which is anticipated to overcome the limitations of traditional vaccines and holds broad development potential (5, 6).

Epitope-focused vaccine design constitutes an effective technical approach to enhancing the efficacy of genetic engineering vaccines (7). This approach has facilitated the development of novel vaccines, including multi-epitope vaccines (8), epitope-chimeric vaccines (9), universal vaccines (10), and dominant epitope-modified vaccines (11). However, few studies have investigated epitope-epitope relationships to date. While immunodominance has been employed to describe relationships between B-cell and T-cell epitopes (12), quantitative research in this area is scarce. We hypothesize that inter-epitope competition on an antigen results in only a small subset of non-overlapping epitopes (distinct antigenic sites) being capable of triggering significant antibody responses in each vaccinated individual. In subsequent sections of this manuscript, non-overlapping epitopes capable of eliciting significant antibody responses will be referred to as “effective epitopes.”

Pasteurella species (predominantly *Pasteurella multocida*) cause pasteurellosis in animals and infections in humans. Pasteurellosis is endemic, leading to animal mortality, impaired animal growth, and reduced feed conversion efficiency—resulting in substantial losses to the livestock and poultry industries (13). *Pasteurella multocida* primarily infects humans via animal bites or scratches, leading to meningitis, sepsis, and other pathologies (14, 15). PlpE is a good protective antigen of *P. multocida*, and its denatured form (i.e., linear epitopes) alone confers sufficient immune protection. In previous work, we identified the non-overlapping linear epitopes of *P. multocida* PlpE and successfully generated PlpE epitope-chimeric protein vaccines (9, 16). Herein, we generated a panel of PlpE epitope-chimeric proteins and determined the number of PlpE effective epitopes in vaccinated individuals.

The gene encoding the PlpE immunodominant region (harboring 6 linear epitopes) was fused (16) to genes encoding protective antigens of other pathogens (Table S1), and inclusion bodies were obtained following expression in *Escherichia coli* engineering strains (Fig. 1A, Fig. S1). The proteins chimerized with PlpE included the P domain of Rabbit Hemorrhagic Disease Virus (RHDV) VP60(9), Bordetella bronchiseptica BcfA (17), and *P. multocida* PtfA(18). Full-length PlpE can be considered a specialized chimeric protein comprising the PlpE immunodominant and non-immunodominant regions.

**Fig. 1.**
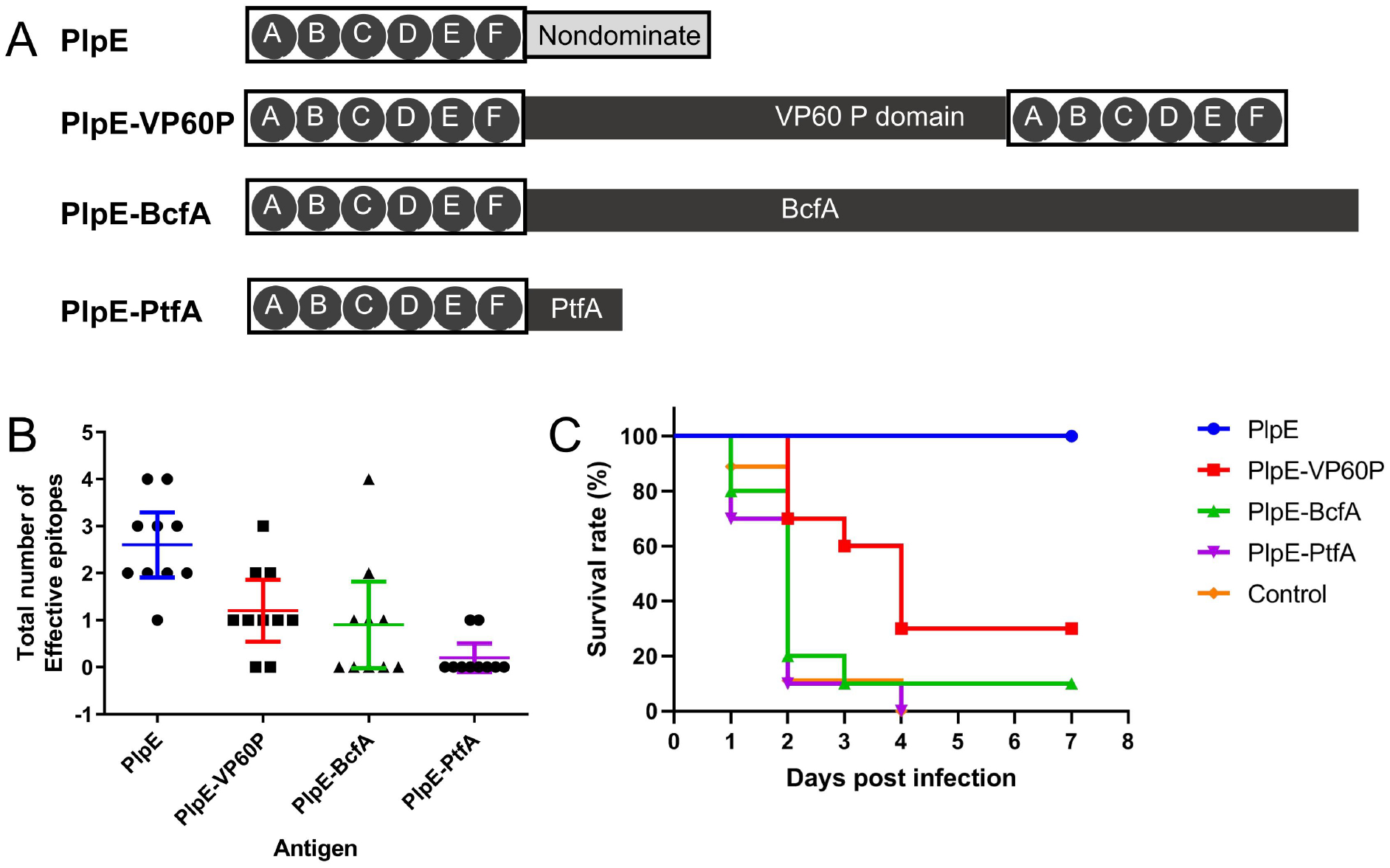
PlpE six-epitope chimeric proteins and their effective epitopes. (A) Schematic of the primary structure of full-length PlpE and its epitope-chimeric proteins. ABCDEF denotes the PlpE immunodominant region, which harbors six linear epitopes (A, B, C, D, E, and F). (B) Number of PlpE effective epitopes in each vaccinated individual. (C) Survival of mice in each antigen-immunized group following challenge with *Pasteurella multocida*.

Female KM mice (4–6 weeks old) were immunized with inclusion bodies solubilized in 8 M urea (to ensure that only linear epitopes trigger immune responses). Complete Freund’s adjuvant was used for primary immunization, and incomplete Freund’s adjuvant for booster immunizations, with a 4-week interval between doses. Serum samples were collected 10 days post-second immunization. Peptide enzyme-linked immunosorbent assay (ELISA) (16) was used to quantify PlpE effective epitopes across vaccinated individuals. The results demonstrated that only a small subset of PlpE epitopes elicited significant antibody responses in each individual (Fig. 1B). The total number of effective epitopes per individual was 2.5 (95% confidence interval (95%CI): 1.909-3.201) for full-length PlpE, and 1.2 (95%CI: 0.5426-1.857), 0.9 (95%CI: 0-1.820), and 0.2 (95%CI: 0-0.5016) for PlpE-VP60P, PlpE-BcfA, and PlpE-PtfA, respectively.

Our results showed that the total number of effective epitopes in PlpE epitope-chimeric proteins did not exceed 3 per vaccinated individual—even though the number of incorporated epitopes in these proteins reached 6. The scaffold protein component of the chimeric protein may itself harbor immunodominant epitopes (19), which compete with PlpE epitopes within the chimeric protein. B cells specific to distinct epitopes compete for T helper (Th) cells and antigens during activation and maturation—leading to mutual interference when these epitopes trigger antibody production (20). This competition results in a lower total number of effective epitopes in PlpE epitope-chimeric proteins—compared to full-length PlpE—in vaccinated individuals.

Immunized mice were challenged via subcutaneous injection with the field virulent *P. multocida* strain X7 (Cps1: L1) (1000 CFU, 40×LD50), and mouse survival is depicted in Fig. 1C. Reduced numbers of effective epitopes impaired the immune protective efficacy of PlpE epitope-chimeric protein antigens, resulting in significantly lower immune protective efficacy of the chimeric epitopes compared to the full-length PlpE group (Log-rank test, P<0.05). The Reduced number of effective epitopes likely underlies the lower efficacy of PlpE chimeric antigen vaccines compared to full-length PlpE reported in the previously work (9).

Beyond the competition between scaffold protein immunodominant epitopes and PlpE epitopes, a competitive relationship also exists between distinct PlpE epitopes—impacting immune efficacy of PlpE chimeric protein. We generated a series of chimeric proteins incorporating subsets of PlpE non-overlapping epitopes. Among these, 4-epitope chimeric proteins (Fig. 2A, Table S2, Fig. S2) were generated herein, whereas 3-epitope, 2-epitope, and 1-epitope chimeric proteins were generated in prior studies (16). Following two immunizations, mouse serum samples were collected to detect antibodies targeting non-overlapping PlpE epitopes.

**Fig. 2.**
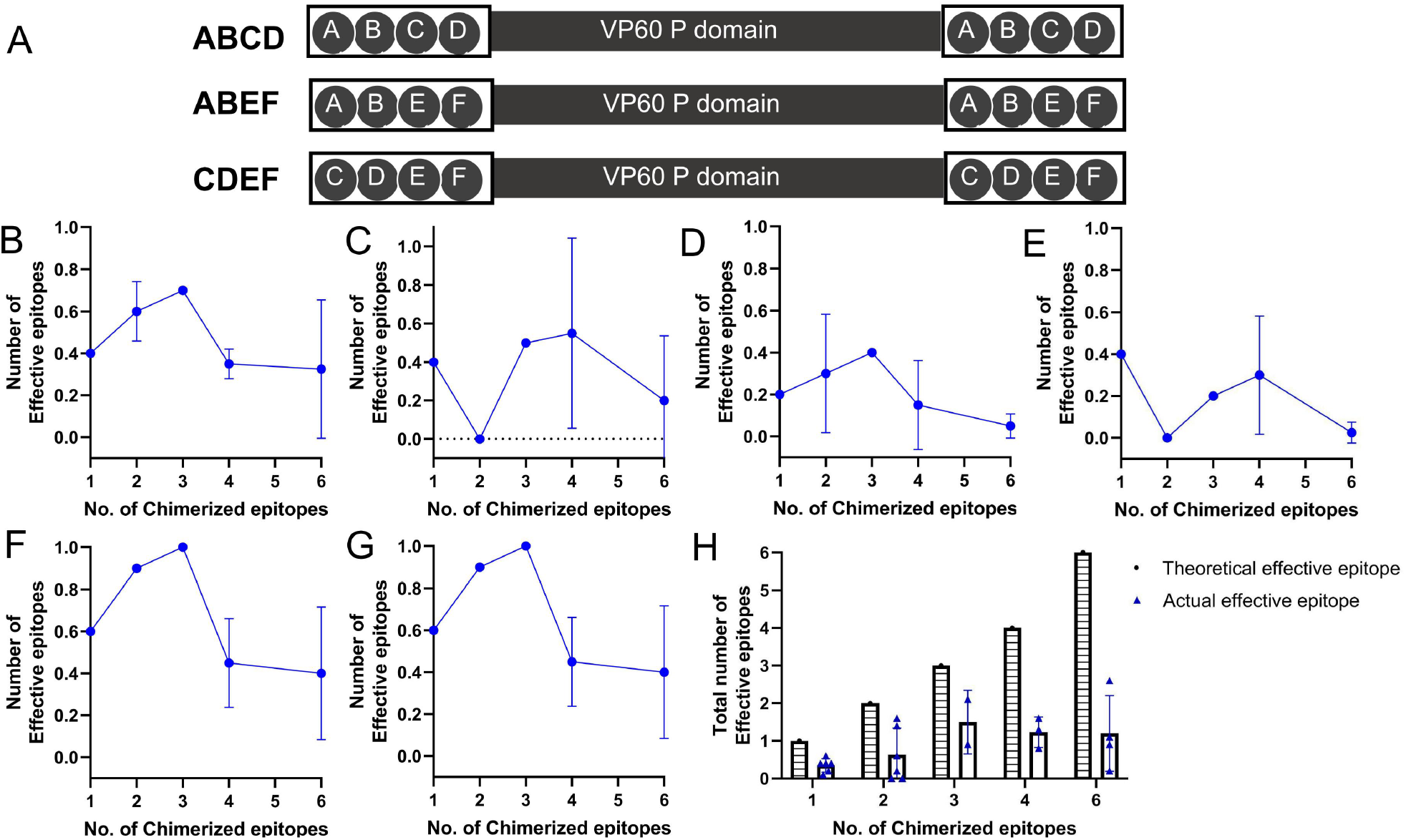
Chimeric proteins with varying numbers of PlpE non-overlapping epitopes and their effective epitopes. (A) 4-PlpE non-overlapping epitope chimeric proteins generated herein. A, B, C, D, E, F denote PlpE epitopes A, B, C, D, E, and F. (B–G) Number changes of effective epitopes of PlpE epitopes in vaccinated individuals as the number of PlpE non-overlapping epitopes in the chimeric protein increased. The effective epitope number = The number of vaccinated individuals in whom the epitope triggers a significant antibody response/the total number of vaccinated individuals. (H) Total number of effective epitopes in proteins chimeric with varying numbers of non-overlapping PlpE epitopes in vaccinated individuals. Total number of effective epitopes = The sum of effective PlpE epitopes contained in the chimeric protein.

With an increase in the number of PlpE epitopes in the chimeric protein, the number of effective epitopes per PlpE epitope (excluding epitope F, which consistently exhibits low immunogenicity) in each vaccinated individual exhibited a trend of initial increase followed by decrease (Fig. 2B–2G, Table S3). Chimeric protein with PlpE epitope A, C, and E exhibited the highest total number of PlpE effective epitopes, 2.1 (95%CI: 1.572-2.628). The initial increase in effective epitopes indicates that a greater number of chimeric epitopes favors competition against scaffold protein epitopes. The subsequent decrease in effective epitopes suggests that inter-epitope competition among PlpE epitopes is intensifying. Ultimately, competition between chimerized epitopes themselves and between chimerized epitopes and scaffold protein epitopes results in the number of PlpE effective epitopes in the chimeric antigen not exceeding 3.

A study investigating B-cell epitopes of virus-like particles demonstrated that while the antigen harbors 10 non-overlapping epitopes, only 2–4 of these epitopes are responsible for generating over 95% of monoclonal antibodies in vaccinated individuals (21). Other studies focused on identifying non-overlapping epitopes via antibodies similarly suggest that the number of effective epitopes of an antigen in an individual is very limited (22). Thus, we propose that the conclusions of this study can be extended to other antigens: in a vaccinated individual, the total number of non-overlapping epitopes that significantly trigger antibody responses is limited (no more than 4, like full-length PlpE). This epitope trait may represent a key factor limiting vaccine efficacy.

To our knowledge, this represents the first report exploring the number limit of effective epitopes in epitope-chimeric proteins in vaccinated individuals—a finding that should be considered in rational vaccine design. The number of exogenous epitopes grafted for constructing novel vaccines should not be excessive, and consideration should be given to the competitive relationships both among the chimerized epitopes themselves and between the chimerized epitopes and the epitopes on the scaffold protein.

## ACKNOWLEDGMENTS

This work was supported by grant C2025204092 from the Hebei Natural Science Foundation.

